# Control of Blood Glucose Level for Type 1 Diabetes Mellitus using Improved Hovorka Equations: Comparison between Clinical and In-Silico Works

**DOI:** 10.1101/2020.10.09.332809

**Authors:** Nur’Amanina Mohd Sohadi, Ayub Md Som, Noor Shafina Mohd Nor, Nur Farhana Mohd Yusof, Sherif Abdulbari Ali, Noor Dyanna Andres Pacana

## Abstract

**Background:** Type 1 diabetes mellitus (T1DM) occurs due to inability of the body to produce sufficient amount of insulin to regulate blood glucose level (BGL) at normoglycemic range between 4.0 to 7.0 mmol/L. Thus, T1DM patients require to do self-monitoring blood glucose (SMBG) via finger pricks and depend on exogenous insulin injection to maintain their BGL which is very painful and exasperating. Ongoing works on artificial pancreas device nowadays focus primarily on a computer algorithm which is programmed into the controller device. This study aims to simulate so-called improved equations from the Hovorka model using actual patients’ data through in-silico works and compare its findings with the clinical works.

**Methods:** The study mainly focuses on computer simulation in MATLAB using improved Hovorka equations in order to control the BGL in T1DM. The improved equations can be found in three subsystems namely; glucose, insulin and insulin action subsystems. CHO intakes were varied during breakfast, lunch and dinner times for three consecutive days. Simulated data are compared with the actual patients’ data from the clinical works.

**Results:** Result revealed that when the patient took 36.0g CHO during breakfast and lunch, the insulin administered was 0.1U/min in order to maintain the blood glucose level (BGL) in the safe range after meal; while during dinner time, 0.083U/min to 0.1 U/min of insulins were administered in order to regulate 45.0g CHO taken during meal. The basal insulin was also injected at 0.066U/min upon waking up time in the early morning. The BGL was able to remain at normal range after each meal during in-silico works compared to clinical works.

**Conclusions:** This study proved that the improved Hovorka equations via in-silico works can be employed to model the effect of meal disruptions on T1DM patients, as it demonstrated better control as compared to the clinical works.

## Introduction

The prevalence of diabetes is steadily increasing all around the world. According to World Health Organization (WHO), the number of people suffered from diabetes has increased from 108 million to 422 million between the year 1980 and 2014[1]. If there is no preventive measures or steps taken in handling this non-communicable disease, it will pose dire consequences towards individual just from having this disease. Diabetes not only affect the lifestyle of patients, but it can also greatly affect their economy due to diabetes-related healthcare expenditure [2]. Diabetes mellitus or commonly known as diabetes is a metabolic disorder characterized by chronic hyperglycaemia or high blood glucose level (BGL) as a result of the defects in insulin secretion, insulin action or both [3]. Insulin is a type of hormone produced by beta cells in the pancreas to regulate the BGL that shall supposedly be in normoglycemic range between 4.0 – 7.0 mmol/L (70 – 120 mg/dL) [4]. Symptoms marked by hyperglycaemia may include increased thirst, frequent urination, extreme fatigue, and blurred vision [5]. Conversely, hypoglycaemia or low blood glucose level, usually occurs due to the overdosing of insulin or blood-glucose lowering medication. Symptoms of hypoglycaemia are such as feeling anxious, confusion, nausea, and in extreme cases the person can slip into coma or death [5].

There are four types of diabetes namely; Type 1 diabetes mellitus (T1DM), Type 2 diabetes mellitus (T2DM), Gestational diabetes (GDM), and other specific types of diabetes^6^. This study focuses on T1DM resulting from autoimmune destruction of pancreatic beta-cells, thus lead to deficient insulin production [3]. As a result, the patients are required to take exogenous insulin injection regularly either via subcutaneously or intravenously in order to maintain their blood glucose level at a normoglycemic range so as to avoid serious complications later on. This phenomenon is known as insulin dependent. Possible diabetes-related complications may include; retinopathy, nephropathy, neuropathy, and cardiovascular disease [6]. Although several factors are associated to have caused T1DM to develop; for instance, due to virus infection, genetics and family history, the actual cause of T1DM is not yet known [3] and there is still no cure found with the present-day knowledge. T1DM is commonly found in white people but less commonly found in Asia due to genetic variations which are more prone to T2DM [7]. T1DM symptoms can be developed and marked rapidly only in a matter of weeks. This type of diabetes is commonly found in children and young adults [7].

The prevalence of T1DM is increasing in many countries including Malaysia particularly in children and adolescents, and this condition will remain even after they turn into adults [8]. Current practice of self-monitoring blood glucose (SMBG) via finger pricks and multiple daily injections (MDI) of insulin can be really painful for T1DM patients. Most T1DM patients use this conventional method to maintain their BGL within safe range (4.0 to 7.0 mmol/L). Thus, this has prompted the doctors and researchers to introduce artificial pancreas device (APD) to T1DM patients in managing their BGL. Although the applications of APD technology in medical and healthcare services have been expanding over the years [9-11], the control algorithms used are still lagging behind. There are few limitations that need to be addressed and one of them involves the uncertainty of delivering the right dosage of insulin into T1DM patient. This is necessary to maintain their BGL to be within normoglycemic range. If the insulin is infused in large amount, the patient may experience hypoglycaemia. If the insulin is infused insufficiently, the patient may experience hyperglycaemia.

An attempt also has been made by previous researchers to improve the existing Hovorka equations [12-15]; however, as of to date, there are still no clinical data used in the simulation work so as to prove the applicability and userability of the newly developed control algorithm using the improved Hovorka equations. Furthermore, a comparative study has to be made between the finding results from the clinical work and in-silico work (simulated result via improved equations) using actual T1DM patients’ data in order to compare their performances in regulating BGL for sustainable purposes in the future. Therefore, the objectives of this study are; 1) to determine the amount of administered exogenous insulin required to regulate the BGL in the normoglycemic range at all times for T1DM patients, 2) to compare the findings between the clinical and in-silico works using actual patient data in terms of its performance, and 3) to determine the userability and applicability of the improved Hovorka equations for model verification.

The study is only limited to the effect of meal disturbances to the BGL without considering other disturbances such as stress and physical activities. Apart from that, hormone used is limited to insulin only and served in regulating the BGL for T1DM. T1DM and its complications have caused substantial effects in the quality of life of the people who suffer from this non-communicable or autoimmune disease. The constant demand of T1DM care such as accurate or healthy eating plan, exercising, keeping track of blood glucose level, and others can be really stressful and painful. The expansion of model equations and algorithm employed in the development of artificial pancreas device in this study will certainly help to decrease the duration and number of clinical trials, hence saving the cost and time for both doctors and patients. Besides that, the artificial pancreas device also helps T1DM patient to live and enjoy a normal life.

## Artificial pancreas device

Artificial pancreas is a device that closely imitates the function of actual pancreas in regulating blood glucose level. In general, artificial pancreas device comprises the following parts; continuous glucose monitoring (CGM) sensor, CGM receiver, control algorithm device (CAD) and continuous subcutaneous insulin infusion (CSII) pump [16]. CGM sensor measures the blood glucose level continuously via the sensor attached on the skin prior to transmitting to the CGM receiver from which it displays the current readings and trends of blood glucose in the form of a graph. The readings are sent to the CAD such as smartphone or personal computer (PC) whereby the algorithms analyse and calculate the insulin doses required. The CAD then interacts with the insulin pump to deliver proper doses of insulin.

### Diabetic Model

Previous works related to managing diabetes had been done especially towards using mathematical modelling in the past 50 years in which these works were developed to simulate the glucose-insulin dynamic system [17-24]. For the purpose of this study, improved Hovorka equations from Hovorka Model [21] are used to facilitate the simulation work. In improved Hovorka equations, additional parameters have been added into the glucose subsystem, insulin action subsystem and plasma insulin subsystem whilst the rest of the equations remain the same [14]. This improvement of the equations is done due to lacks of interaction between the parameters and variables in the insulin action subsystem and mass of glucose in accessible compartment (Q_1_) and non-accessible compartment (Q_2_) [14]. The results had caused a change in behaviour in Q_1_ and Q_2_ when compared to Hovorka model [21]. Therefore, the improved Hovorka equations are expected to be more precise and highly efficient in stabilizing glucose-insulin regulatory system for T1DM. More details on the improved Hovorka equations can be found in the methodology section.

### Control Systems

In order for the artificial pancreas to function accordingly, certain control algorithms need to be implemented. There are varieties of control strategies used to control the BGL in T1DM patients. The control strategies proposed for the artificial pancreas are namely; proportional-integral-derivatives (PID) control, neural network control, fuzzy logic control, and model predictive control (MPC) as can be encountered in the literature [25–29]. This study employs enhanced model predictive control (eMPC). The enhanced model predictive control (eMPC) or multi-parametric model predictive control (mp-MPC) possesses advantages in which the on-line optimisation can be done via off-line optimisation allowing for better implementation of MPC [30]. Unlike MPC; it requires on-line optimisation, thus slower the systems. The application of mp-MPC can be seen in the area where MPC is not capable of penetrating systems involving fast dynamics or fast sampling time or where the cost and size of the controller dominate the selection of hardware [30]. Some of the applications of mp-MPC include automotive (catalytic converters), biomedical (artificial organs and drug delivery) and industrial (robotics and process control) [30]. Past work using eMPC as control strategy can be seen in the literature [31].

## Methodology

### Data collection and extraction

Prior to performing this study, written approval to collect patient’s data must first be obtained from a UiTM Research Ethics Committee. Data collection from the actual T1DM patients are required for clinical works and subsequent interviews conducted during the patient’s appointment with a paediatrician at UiTM Medical Specialist Centre, Sungai Buloh, Selangor with the consent of the parents. Data collected include the patients’ social demography, which are namely; name, date of birth, gender, race, body weight, height, body mass index, and year diagnosed with T1DM, type of meal intake from breakfast, lunch and dinner, amount of CHO taken every meal, pre-meal BGL via fingerpicks and the amount of insulin administered before meal. Data collected during the appointment day are termed as clinical data in this study and all the data were taken from Patient 1 throughout Day 1 to Day 3 observations. These clinical data were subsequently fitted into the improved Hovorka equations for simulation purposes using MATLAB in the in-silo works. Both results from clinical and in-silico works were then compared, accordingly. Figure 1 shows the schematic flow diagram of the methodology, and Table 1 shows brief information of patient 1.

**FIG.1.**
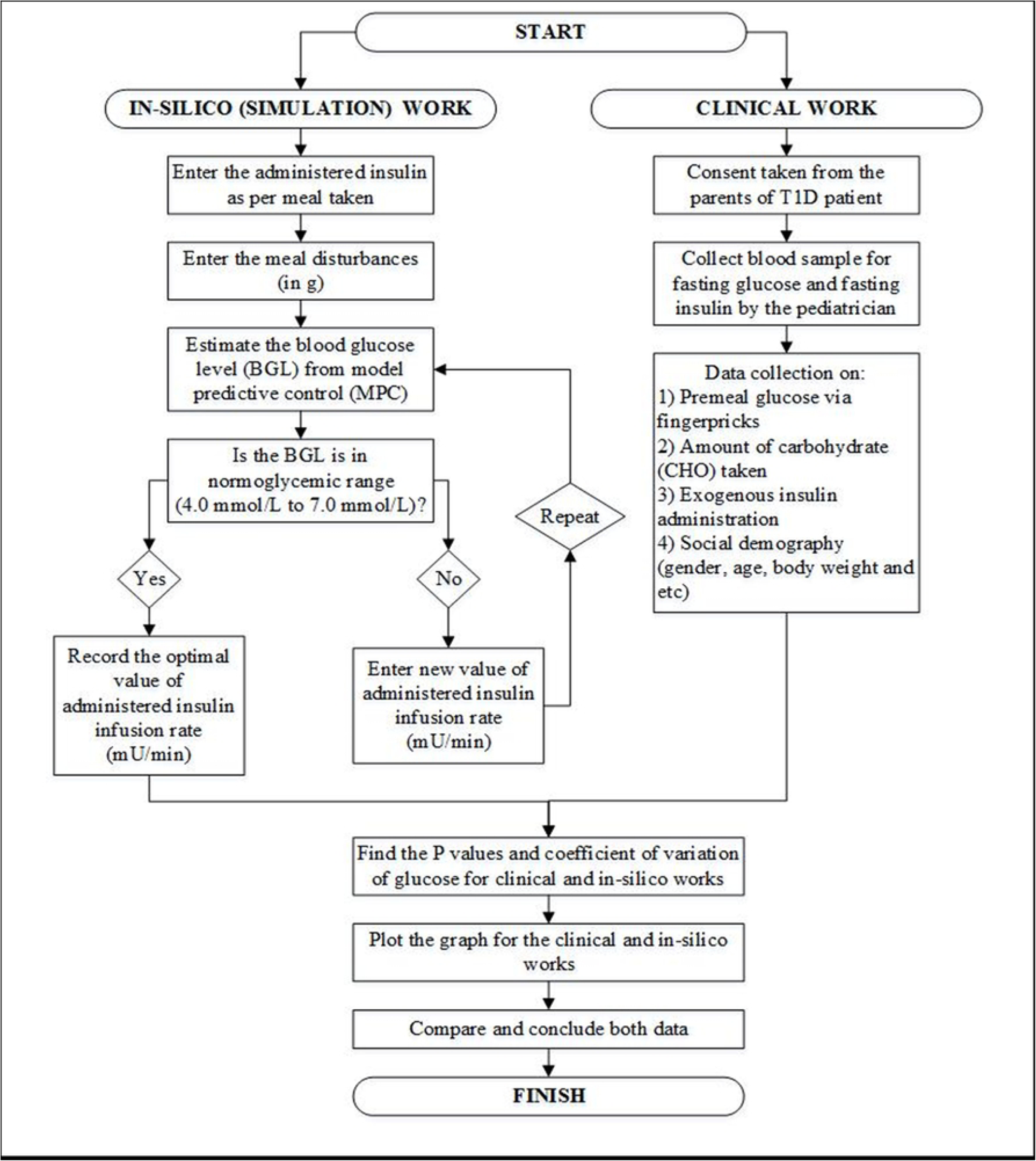
Schematic Flow Diagram of the Methodology.

### In-Silico (simulation) works using Improved Hovorka Equations

As mentioned above, data from the clinical works were used to simulate the improved Hovorka equations in the in-silico works. The improved Hovorka equations were adopted from Yusof et al.[14] The glucose subsystem, plasma insulin subsystem and insulin action subsystem equations in the original Hovorka method have been modified while other formulas remain unchanged. Figure 2 shows the schematic flow diagram of the improved equations adapted from [24]. The improved equations are described as follows:

**FIG. 2.**
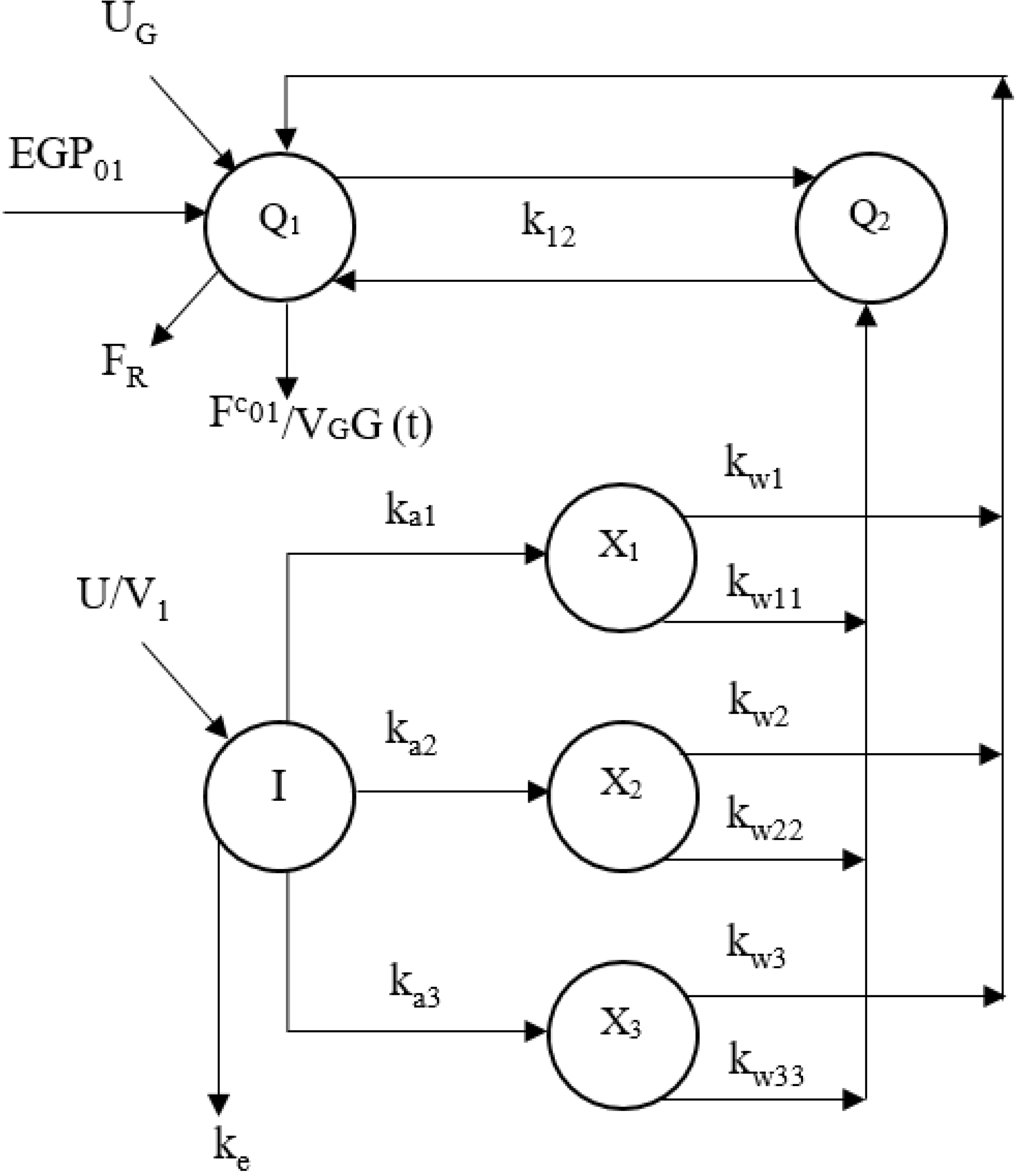
Schematic flow diagram of lmproved.

### Glucose subsystem

In the improved Hovorka equations, the glucose subsystem can be represented by the following equations:

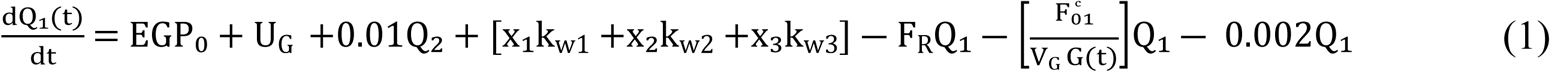

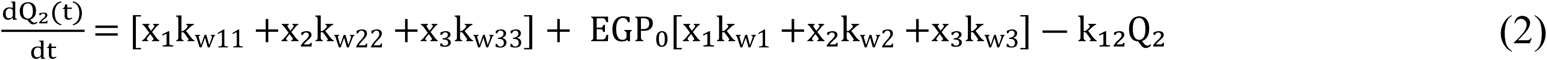

Where the total non-insulin dependent glucose flux, F_01_^c^ (mmol/min), and renal glucose clearance, F_R_ (mmol/min), can be found as in equations (3) and (4).

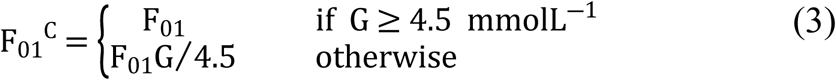

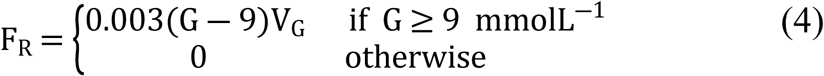

Where,

Q_1_(t) = mass of glucose in accessible compartment (mmol)

Q_2_(t) = mass of glucose in non-accessible compartments (mmol)

U_G_ = gut absorption rate (mmol/min)

Equation for meal disturbances is represented by equation (5)

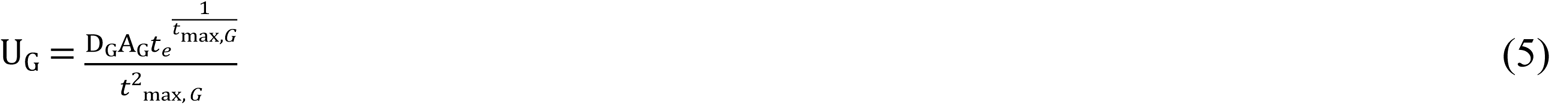

Where,

D_G_ = amount of carbohydrate (CHO) digested (mmol)

A_G_ = carbohydrate bioavailability (unitless)

t_max,G_ = time-to-maximum of CHO absorption (min)

### Plasma insulin subsystem

The insulin subsystem can be represented by the following equations:

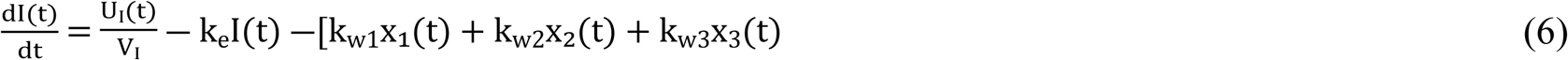

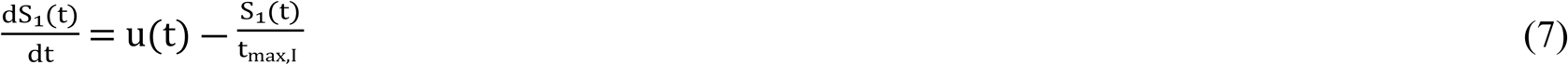

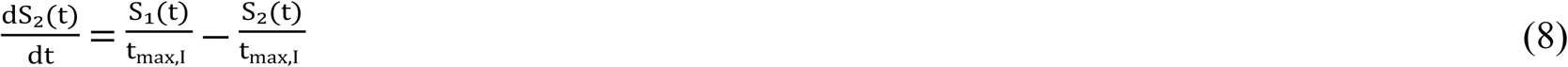

Where,

S_1_ = insulin sensitivity in the accessible compartment (mU)

S_2_ = insulin sensitivity in the non-accessible compartment (mU)

U_I_ = insulin absorption rate (mU/min)

U(t) = exogenous insulin input (mU/min)

t_max,I_ = time-to-maximum of insulin absorption (min)

In the improved equations, few variables are added to the plasma insulin concentration equation as seen in (6).

### Insulin action subsystem

Equations (9) to (11) represent the insulin action subsystem as follows:

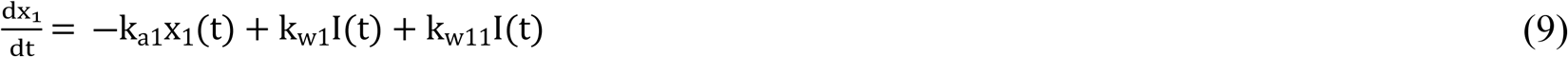

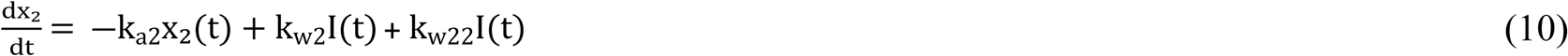

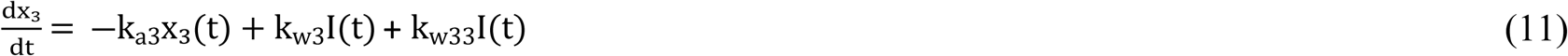

Constants and parameters involved in the equations are shown in the Tables 2 and 3.

The simulation work on the meal disturbances was based on the actual patients’ daily meals intake. Table 4 shows an example of amount of daily meals intake specifically carbohydrate (CHO) suggested for a male patient aged 13 years old (Patient 1). The suggested meal intake is based on total daily calorie requirement which varies according to age and gender as shown in Table 5. It consists of breakfast, lunch and dinner at which this information is adopted into the simulation work using the improved Hovorka equations. Generally, 50% of calorie intake comes from CHO [30]. The calculations of CHO intake are shown in the following session.

### Sample calculation of CHO intake for T1DM patient

Sample calculations of CHO intake for the T1DM male patient aged 13 years old with body weight of 27 kg (Patient 1) are shown as follows. The daily energy (calorie) requirement by weight is based on Table 5 for a male patient aged between 11 to 14 years old [33]. This serves as a guideline for T1DM patient.

Daily energy (calorie) requirement by weight = 55 kcal/kg/day

So,

55 kcal/kg/day × 27 kg = 1485 kcal/day

Therefore,

1485/2 = 742.5 kcal/day CHO

Convert the unit whereby 4 kcal = 1g CHO

742.5/4 = 185.6g CHO

The bolus (exogenous) insulin required per day can be calculated based on the amount of CHO intake, i.e. 117g CHO. In general, 1-unit of rapid acting insulin can cover 10 to 15 grams of carbohydrate [34]. However, this range can vary from 6 to 30 grams depending on the individual’s sensitivity to insulin as the insulin sensitivity for individual may vary for different period of time [34]. The total daily dose (TDD) of insulin can be calculated based on the insulin-to-carbohydrate ratio (ICR) used which is 1:15 as shown in equation (12).

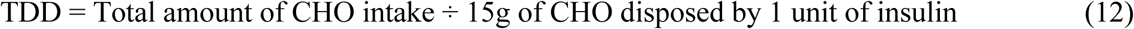

Therefore,

TDD = 117g CHO ÷ 15g of CHO disposed by 1 unit of insulin

TDD = 7.8 ≈ 8 units of rapid acting insulin

From the calculation above, the total daily dose of insulin required to cover the 117g of CHO intake is 8 units.

The enhanced Model Predictive Control (eMPC) utilized in this study was to regulate future glucose concentrations based on the system model. The model was a nonlinear approximation of the system in which measurement for future glucose concentration was predicted based on the previous inputs namely; insulin infusion rate and meal disturbances. Constraint was imposed such that the blood glucose level, G(t), was set at normoglycemic range (4.0 – 7.0 mmol/L) and the insulin infusion rate, u(t), was between (0 – 100 mU/min) based on the current insulin pump specification [21-22].

Initial values for S_1_, S_2_, x_1_, x_2_, and x_3_ were set at zero since the insulin had not been injected into the patient’s body and the insulin being administered which caused glucose transport/distribution, glucose disposal and EGP had not yet occurred [24]. The bolus insulin for each meal was obtained from trial and error basis in order to minimize the BGL to normal range while not experiencing hypoglycaemia (≤4.0 mmol/L). The time at which insulin was administered should be between 30 minutes before meal and at meal time.

## Results and Discussion

### Comparison of results between Clinical and in-silico works for Patient 1 on Day 1

Figure 3 shows the graph of blood glucose level (BGL) versus time for clinical and simulation works for 24-hours of patient 1 on day 1. The blue and yellow lines represent the clinical and simulation data, respectively, while the red lines represent the target range of BGL within 4.0 to 7.0 mmol/L (normoglycemic range). In the clinical work, the patient needed to do self-monitoring blood glucose (SMBG) via finger prick before and 2-hours after every meal and multiple daily injections (MDI) of insulin to maintain his BGL in normoglycemic range. On the other hand, as the patient woke up in the morning and started to consume meals on Day 1, his BGL was measured and monitored continuously and the exogeneous insulin was infused automatically (by a micro pump as set and programmed in the control algorithm) throughout the day in the in-silico works. The patient had three meals per day which were taken at the same time for three consecutive days; namely breakfast (7.30 am), lunch (1.30 pm) and dinner (7.00 pm).

**FIG. 3.**
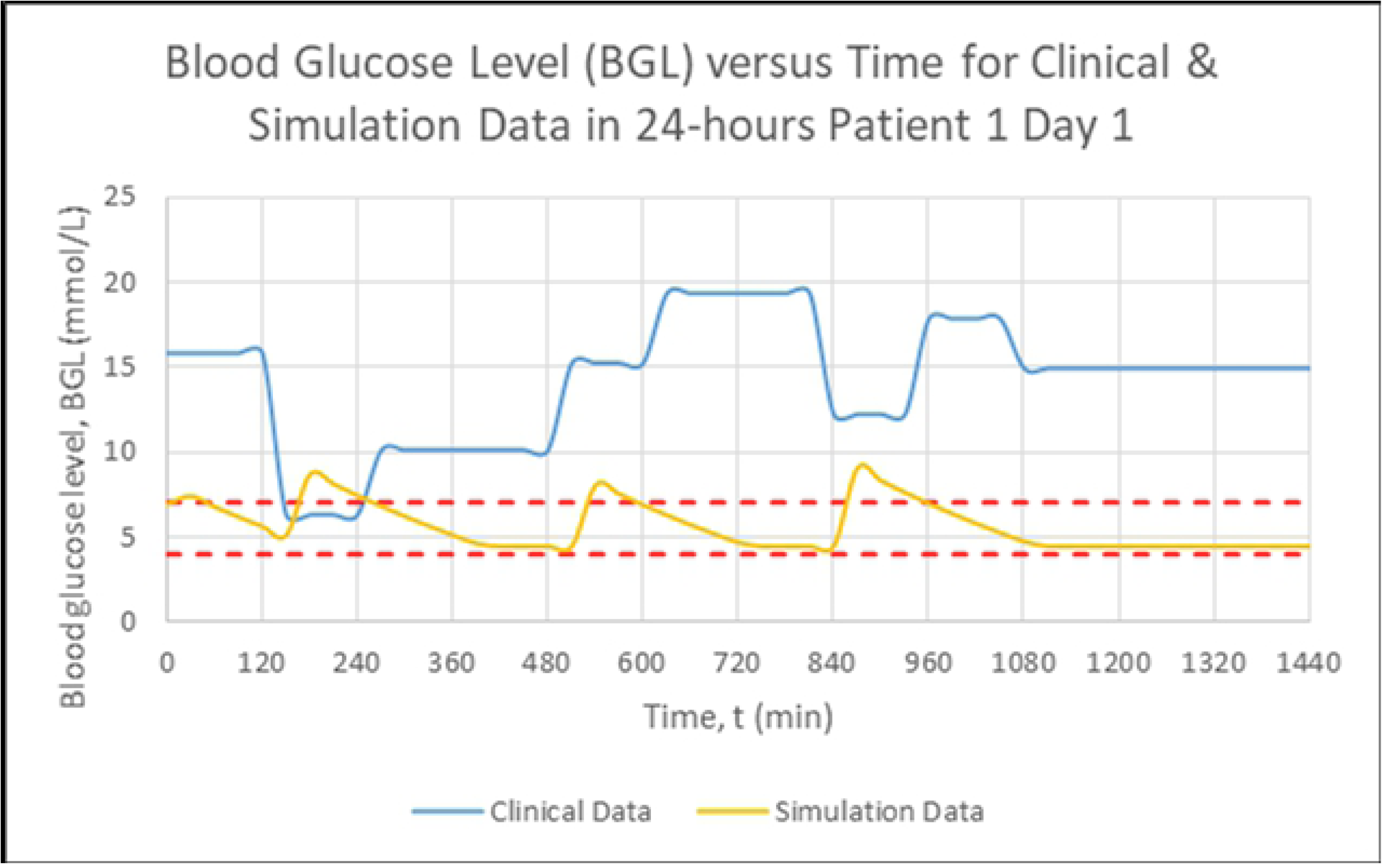
Blood Glucose Level (mmol/L) versus Time.

Based on the graph for clinical works (blue line) as shown in Figure 3, patient 1 has been experiencing hyperglycaemia for most of the time within 24-hours. Patient 1 managed to achieve normoglycemic range during breakfast time (7.30 am) only within t = 150 to 240 min. The lowest BGL recorded was 6.30 mmol/L during this time. Peak BGL recorded was 19.30 mmol/L at 2-hours post lunchtime (3.30 pm) within t = 630 to 810 mins. For the simulation works (yellow line), the patient was able to achieve normoglycemic range in a significant time of the day with the lowest BGL recorded was 4.50 mmol/L and finally stabilized in the end, starting at t = 1110 min (11.30 pm). Although there is a fluctuation of BGL after every meal, which is normal, the BGL does not rise drastically to a harmful level. Peak BGL recorded was 9.13 mmol/L during dinner and was close to the normoglycemic range.

Figure 4 shows the simulation work of BGL versus time for 24-hours of patient 1 on day 1. The simulation work was done in MATLAB R2016a using data from the clinical work as previously mentioned. The simulation work started at 5.00 am (t = 0 min), for which basal insulin was administered at 66.67 mU/min. Bolus insulin was administered 30 minutes before meal time. At t = 120 min, the first bolus insulin of 100 mU/min was administered before breakfast at 7.30 am (t = 150 min). During breakfast, the patient consumed 36 g of CHO which caused the BGL to increase rapidly from 5.16 to 8.77 mmol/L within 30 minutes of meal duration. The BGL gradually decreased over time until it reached 4.5 mmol/L and stabilized. At t = 480 min, second bolus insulin was administered at 100 mU/min. The patient consumed 36 g CHO at lunchtime (1.30 pm) with 30 minutes meal duration. Shortly afterwards, the BGL increased from 4.50 to 8.16 mmol/L. The final bolus insulin of the day was administered at t = 810 min at the exogeneous insulin infusion rate of 100 mU/min. Patient 1 had a dinner at 7.00 pm (t = 840 min) for half an hour and consumed 45 g CHO. The BGL rose from 4.50 to 9.13 mmol/min after 30 minutes of the meal time, the highest BGL recorded. After around one and a half hours (1 ½ hour), the BGL reached the normoglycemic range of 6.93 mmol/L at t = 965 min. Figures 5 to 7 show the detailed day 1 simulation work profiles of BGL versus time for breakfast, lunch and dinner, respectively.

**FIG.4.**
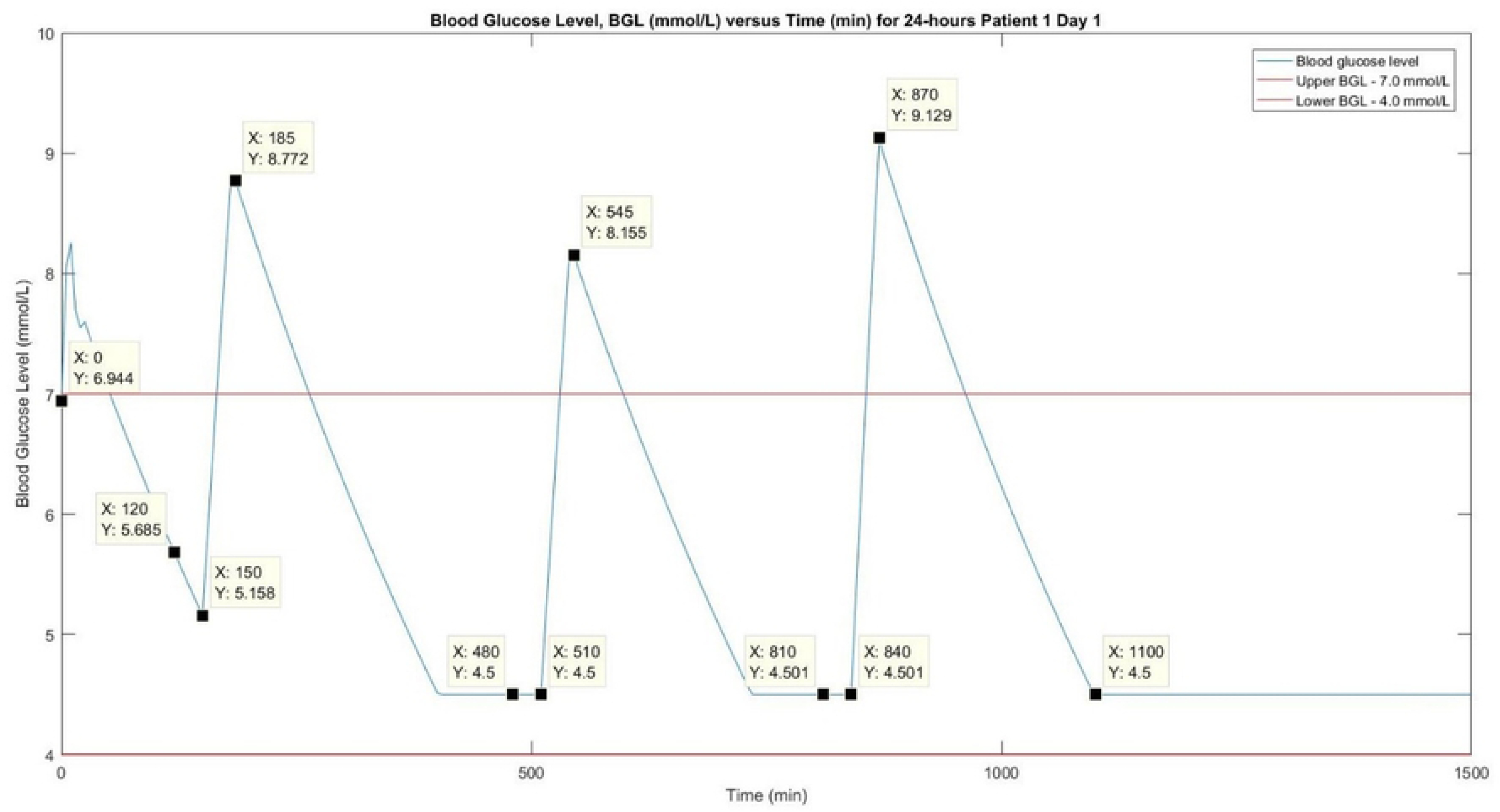
Simulation Works of Blood Glucose Level (mmol/L) versus.

**FIG.5.**
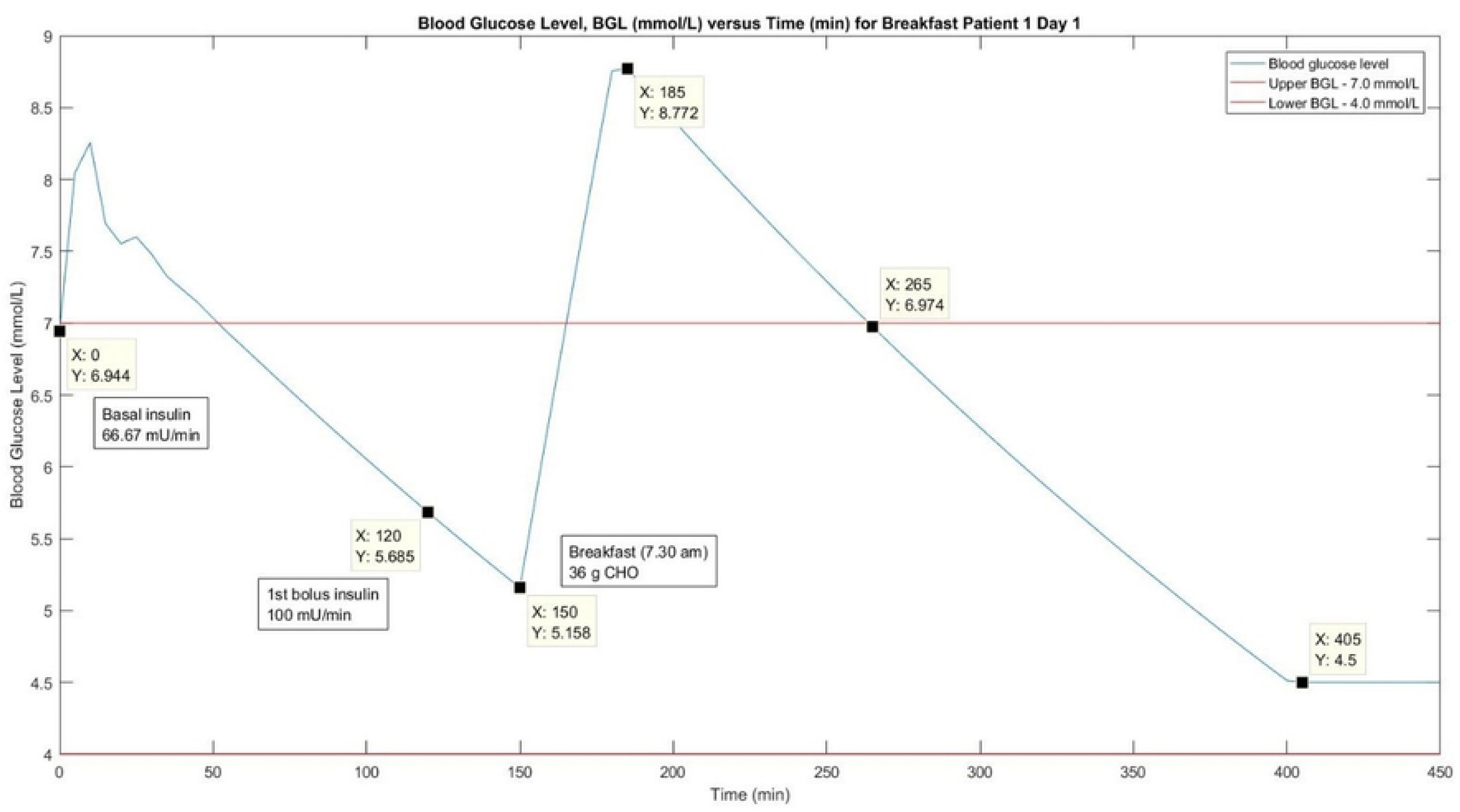
Simulation Works of Blood Glucose Level (mmol/L) versus.

**FIG.6.**
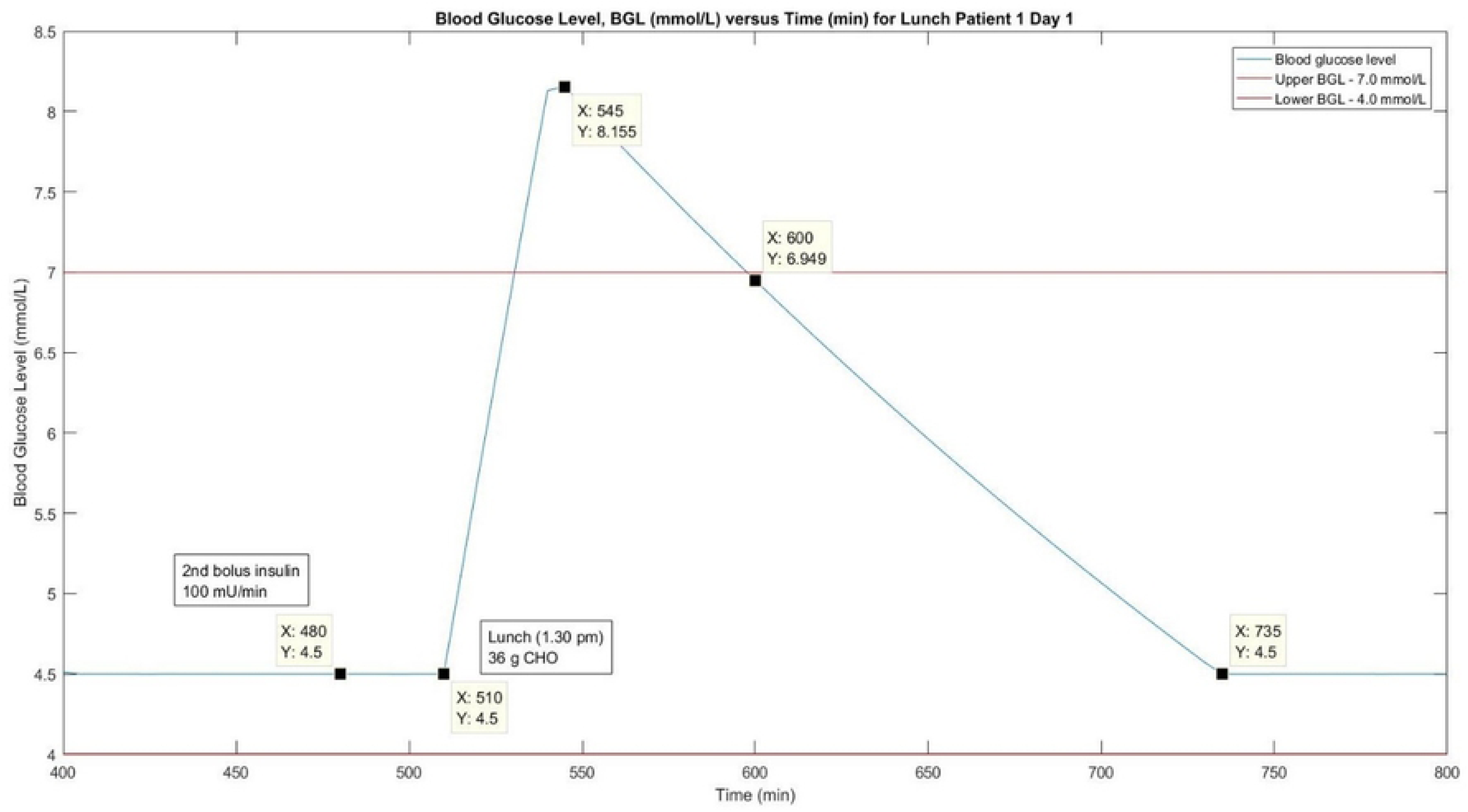
Simulation Works of Blood Glucose Level (mmol/L) versus.

**FIG. 7.**
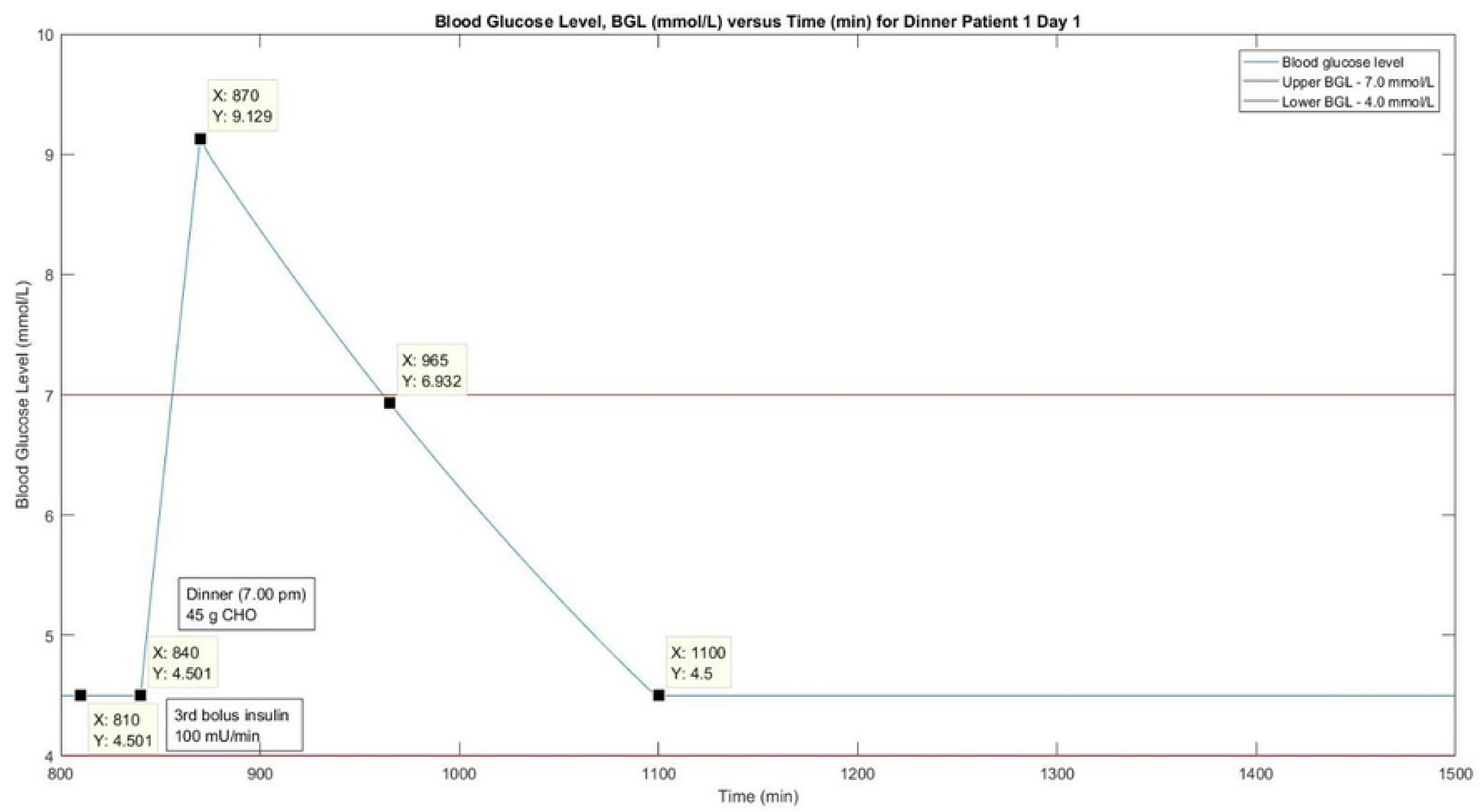
Simulation Work of Blood Glucose Level (mmol/L) versus.

Table 6 shows the comparison of BGL (at peak) between clinical and simulation works for patient 1 on day 1. The BGL for clinical work was significantly higher than the simulation work. The clinical work recorded highest BGL during lunch at 19.30 mmol/L, while the simulation work showed at 9.13 mmol/L during dinner. This result indicates that simulation work has indeed better control of BGL over clinical work. Tables 7 and 8 show the analysis of clinical and simulation data for Patient 1 on Day 1, respectively. Based on both tables, p values were recorded below 0.05 which indicated that all the clinical and simulation data were statistically acceptable and in good order.

### Comparison of results between Clinical and in-silico works for Patient 1 on Day 2

Figure 8 shows the graph of clinical and simulation works for 24-hours of the patient 1 on day 2. As similar to the day 1, the patient has experienced hyperglycaemia for most of the time within 24-hours in the clinical work (blue line). Patient 1 managed to achieve normoglycemic range during breakfast time (7.30 am) only within t = 150 to 240 min, and the BGL recorded was 5.00 mmol/L. Peak BGL was recorded relatively high at 30.00 mmol/L within which the time, t = 270 to 600 min. The high BGL recorded was due to a high amount of carbohydrate (CHO) consumed on that particular day. For the simulation work (yellow line), the patient was able to achieve normoglycemic range most of the time in 24-hours with the lowest BGL recorded was 4.50 mmol/L and stabilized afterwards starting at t = 810 min (6.30 pm). In comparison to the day 1, BGL started to maintain late at night at t = 1110 min (11.30 pm). Skipping dinner mainly affects the BGL to achieve normoglycemic range faster in day 2. Peak BGL recorded was 11.01 mmol/L during lunch.

**FIG.8.**
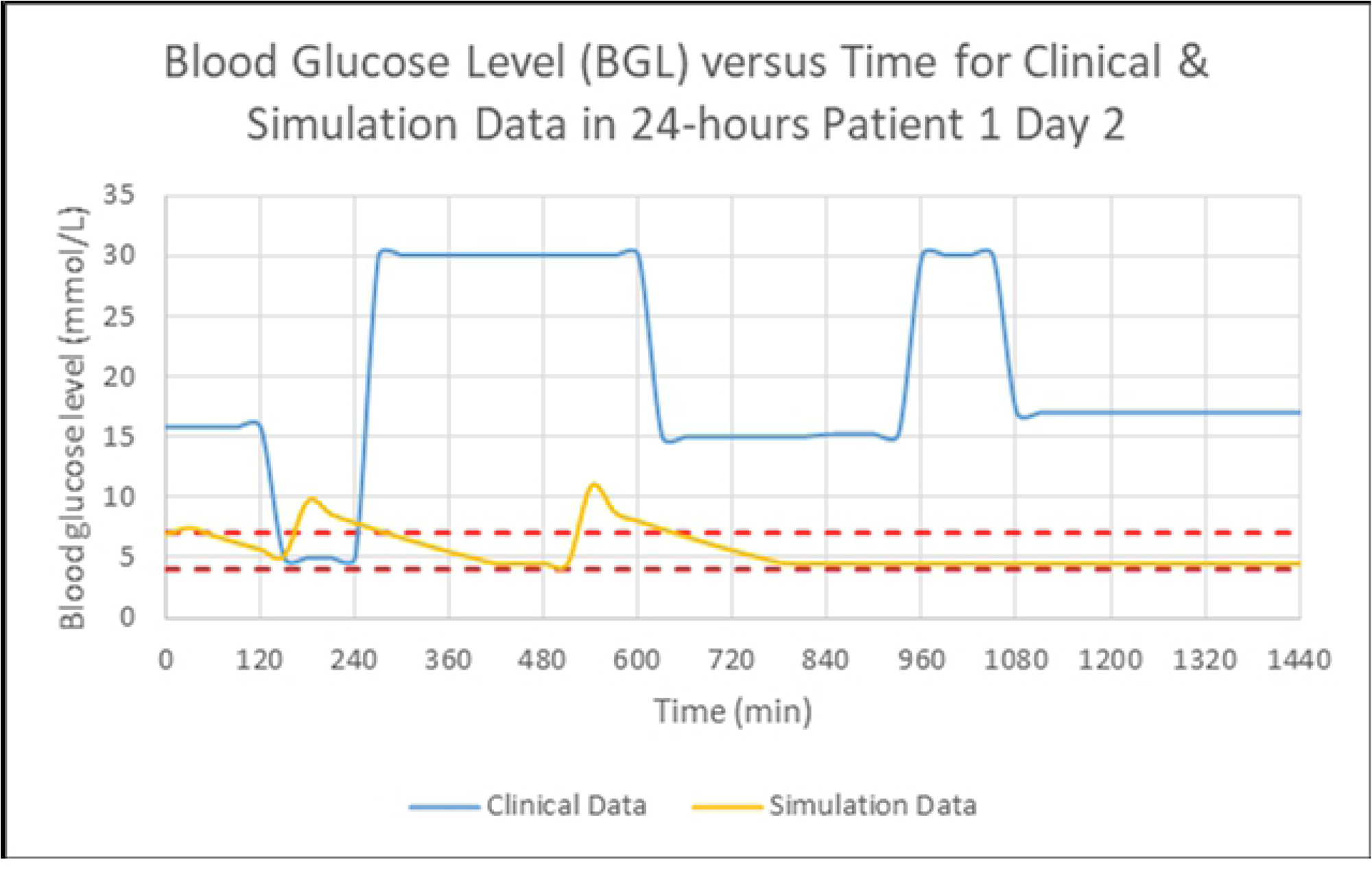
Blood Glucose Level (mmol/L) versus Time.

Figure 9 shows the simulation work of BGL versus time for 24-hours of patient 1 on day 2. The simulation work started at 5.00 am (t = 0 min) in which basal insulin was administered at 66.67 mU/min. Bolus insulin was administered 30 minutes before meal time. At t = 120 min, the first bolus insulin of 100 mU/min was administered before breakfast at 7.30 am (t = 150 min). During breakfast which took approximately 30 minutes to finish, the patient consumed 48 g of CHO. The BGL recorded an increase from 5.16 to 9.74 mmol/L within that period. After t = 180 min, the BGL slowly decreased until it reached 4.5 mmol/L before the next meal. At t = 480 min, second bolus insulin was administered at 100 mU/min. The patient had a lunch at 1.30 pm for approximately 30 minutes and consumed 88 g CHO. The BGL later increased from 4.50 to 11.01 mmol/L, the highest BGL ever recorded on the day 2. Despite not having dinner, the final bolus insulin was administered at t = 810 min at 100 mU/min to regulate BGL within a safe range. At t = 650 min, the BGL managed to reach the normoglycemic range of 6.95 mmol/L. Figures 10 and 11 show the detailed simulation work profiles of BGL versus time for breakfast and lunch for day 2, respectively.

**FIG.9.**
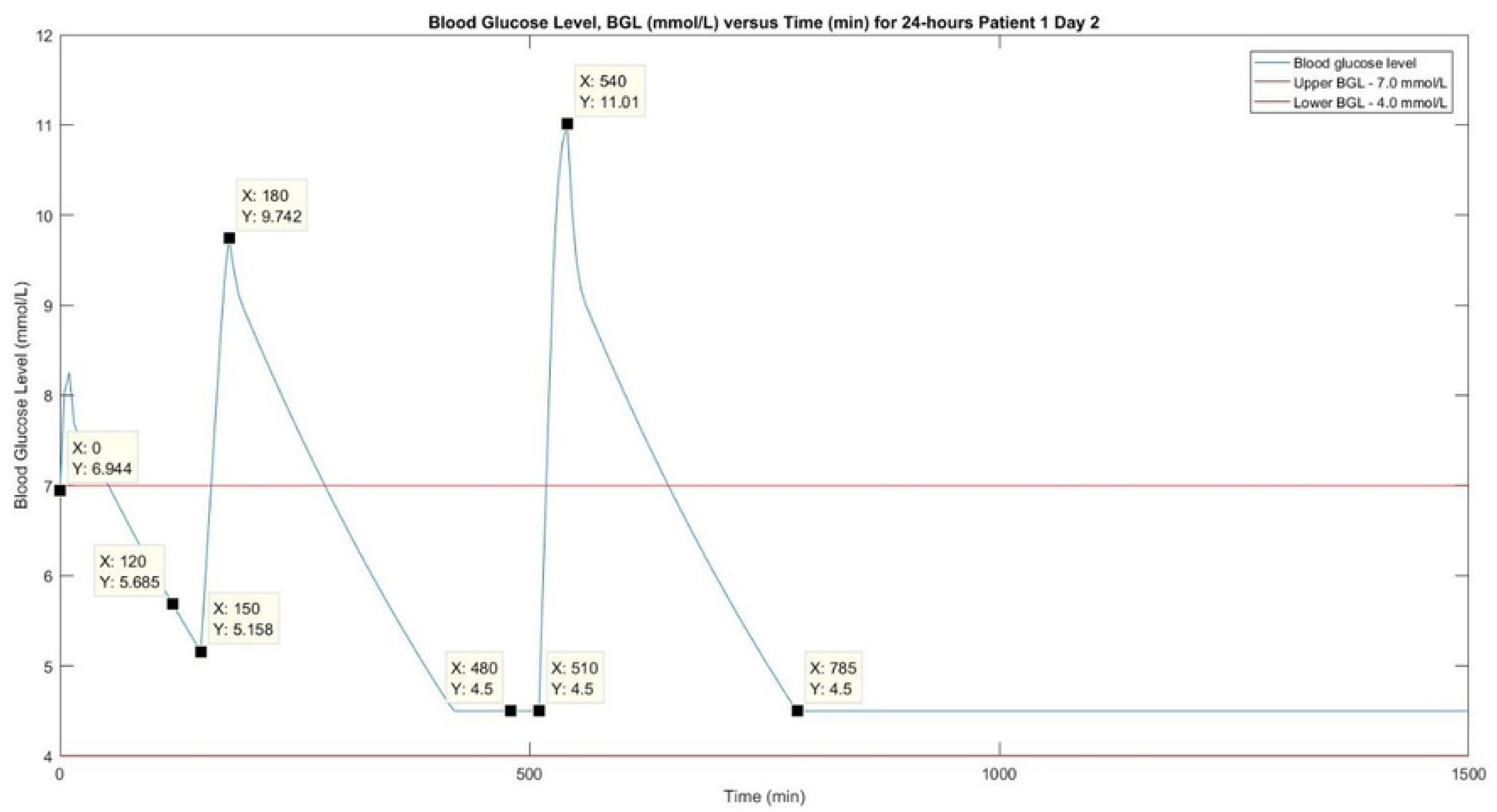
Simulation Work of Blood Glucose Level (mmol/L) versus Time.

**FIG. 10.**
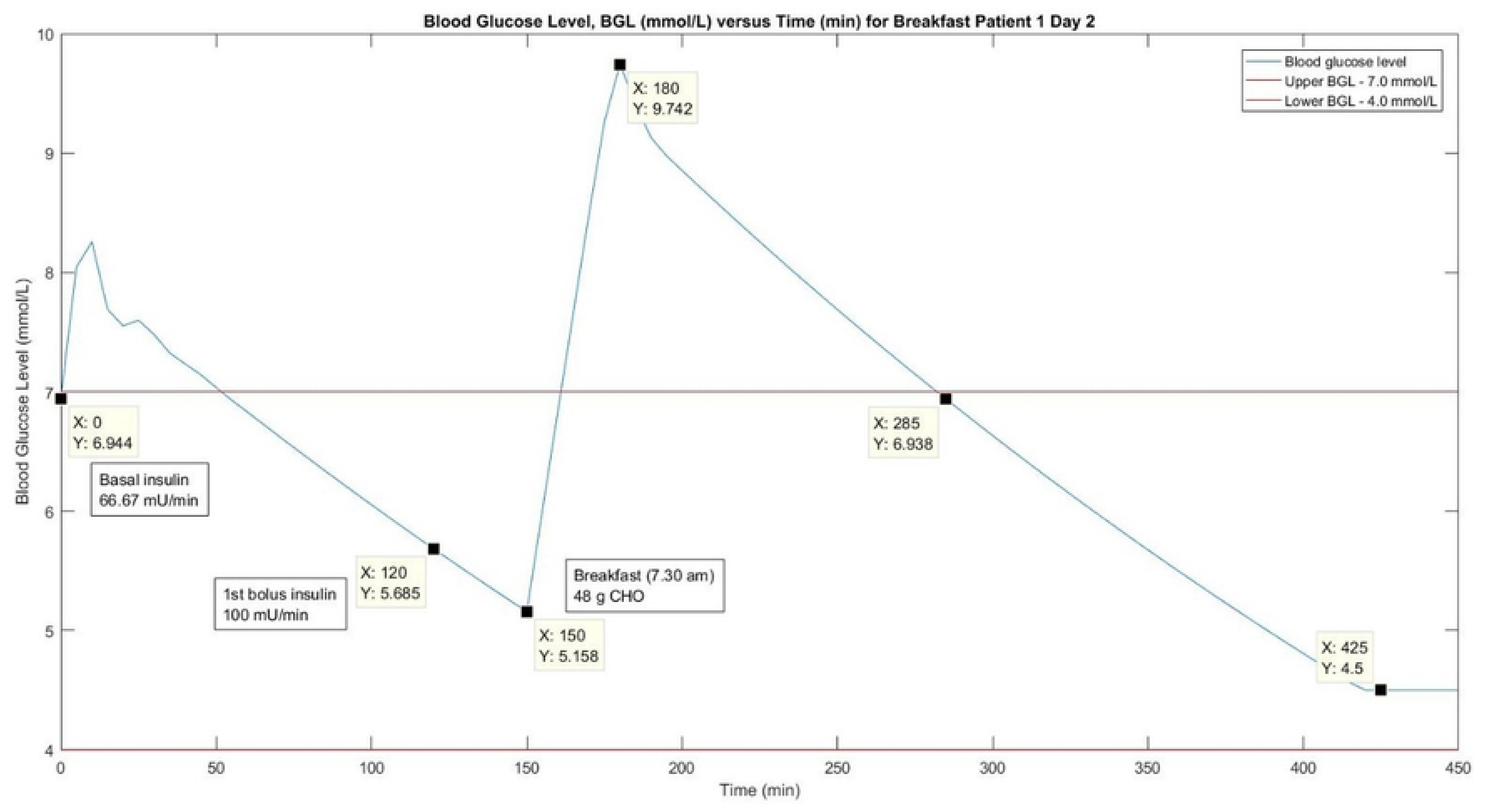
Simulation Work of Blood Glucose Level (mmol/L) versus.

**FIG. 11.**
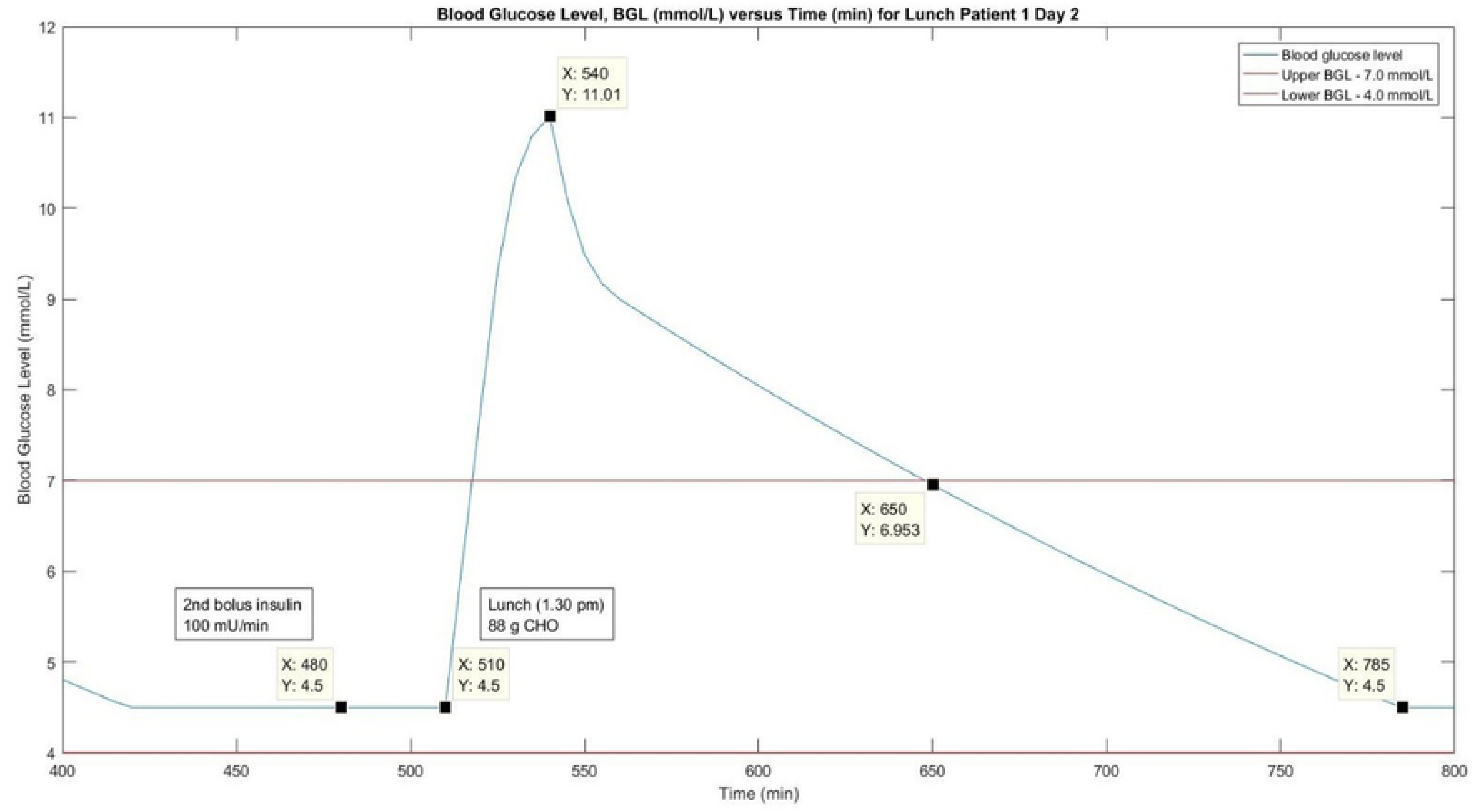
Simulation Work of Blood Glucose Level (mmol/L) versus.

Table 9 shows the comparison of BGL at peak condition between clinical and simulation works on day 2. It demonstrated the same condition as in day 1 in which clinical work recorded significantly high BGL at 30.00 mmol/L during breakfast and lunch compared to the simulation work at 11.01 mmol/L during lunch only. Similarly, it was proven that the simulation work managed to control BGL better than in the clinical work. Table 10 shows the clinical data analysis of Patient 1 on day 2. Based on the table, p value for x-variable 1 was recorded higher than 0.05 which indicated that that the data was statistically not strong and reliable. This might be due to unavailability of data for meal time during dinner as the patient skipped his dinner on day 2. However, Table 11 shows that p values were below 0.05 for the simulation data. This again proved that the simulation work yielded much reliable, strong data which, in turn, resulted in better control of the patient’s BGL.

### Comparison of results between Clinical and in-silico works for Patient 1 on Day 3

Figure 12 shows the profile of BGL versus time for clinical work (blue line) and simulation work (yellow line) for 24-hours of patient 1 on day 3. Based on Figure 12, patient 1 frequently experienced hyperglycaemia episode within 24-hours during clinical works, and the trend was similar to the previous two days. Patient 1’s BGL was recorded at 4.30 mmol/L during breakfast time (7.30 am) at t = 150 to 240 min, which was close to hypoglycaemia. Nevertheless, his BGL reached its peak at 17.40 mmol/L within t = 270 to 510 min right after he consumed his breakfast. For the simulation work, the patient’s BGL remained at the normoglycemic range most of the time. The highest BGL ever recorded was 9.14 mmol/L. After dinner (7.00 pm), patient’s 1 BGL gradually stabilized at 4.50 mmol/L at t = 1110 min (11.30 pm) and thereafter.

**FIG. 12.**
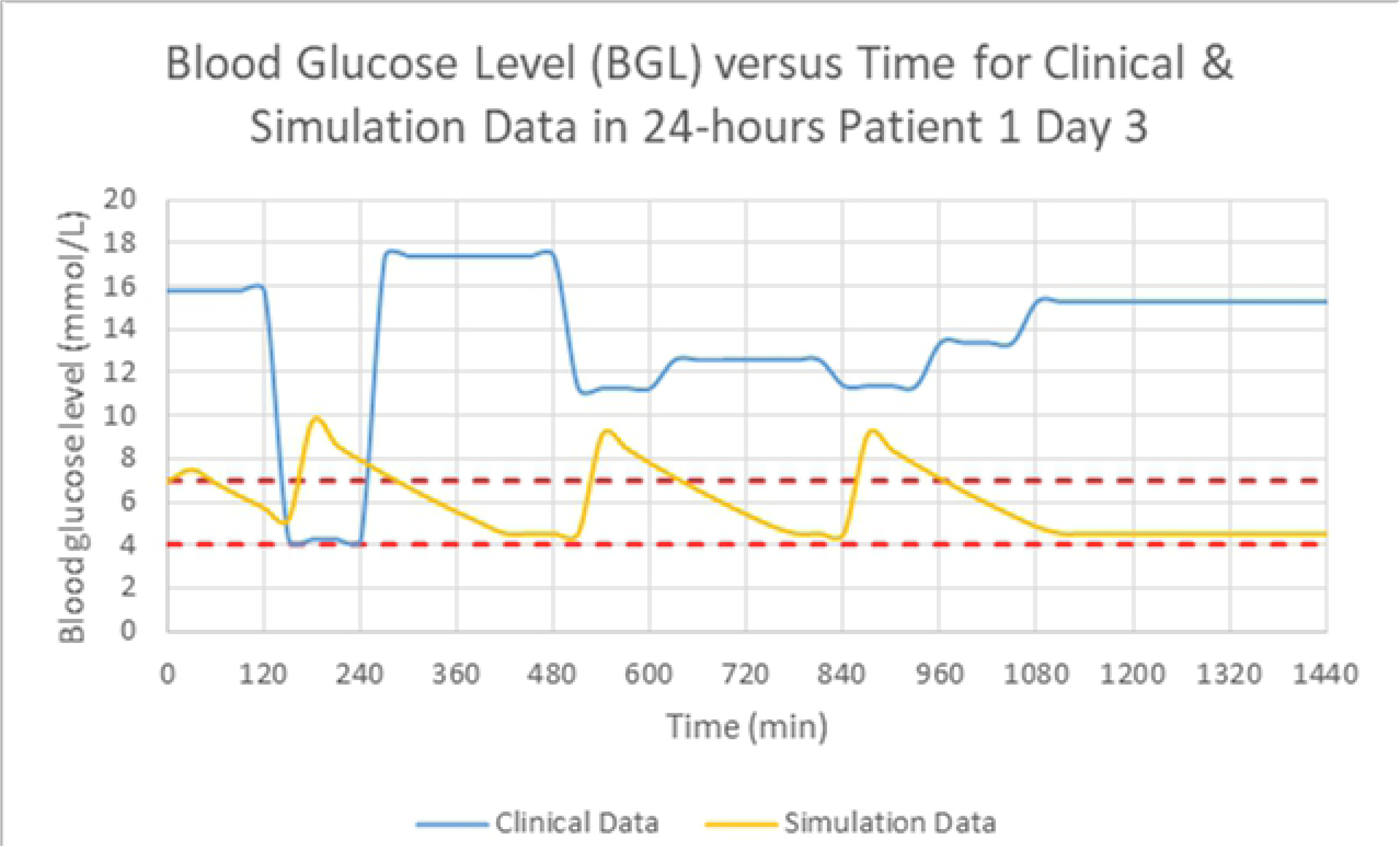
Blood Glucose Level (mmol/L) versus Time.

Figure 13 shows the simulation work profile of BGL versus time for 24-hours of patient 1 on day 3. Similarly, the simulation started at 5.00 am (t = 0 min) in which basal insulin was administered at 66.67 mU/min. Bolus insulin was administered 30 minutes before mealtime. At t = 120 min, the first bolus insulin of 100 mU/min was administered before breakfast at 7.30 am (t = 150 min). During breakfast, the patient consumed 48 g of CHO. The BGL recorded an increase from 5.16 to 9.74 mmol/L at t = 180 min. It slowly decreased thereafter until it reached normoglycemic range at t = 285 min, after 1 hour and 45 minutes. At t = 480 min, second bolus insulin was administered at 100 mU/min. The patient consumed 45 g CHO during lunch (1.30 pm) for approximately 30 minutes. The BGL later increased from 4.50 to 9.13 mmol/L. The final bolus insulin was then administered at 83.33 mU/min at t = 810 min. When the patient consumed 45g CHO during dinner at t = 840 min (7.00 pm), the BGL recorded an increase from 4.50 to 9.14 mmol/L prior to gradually descending upon reaching the normoglycemic range at t = 1100 min. Figures 14 to 16 show the detailed day 3 simulation profiles of BGL versus time for breakfast, lunch, and dinner, respectively.

**FIG. 13.**
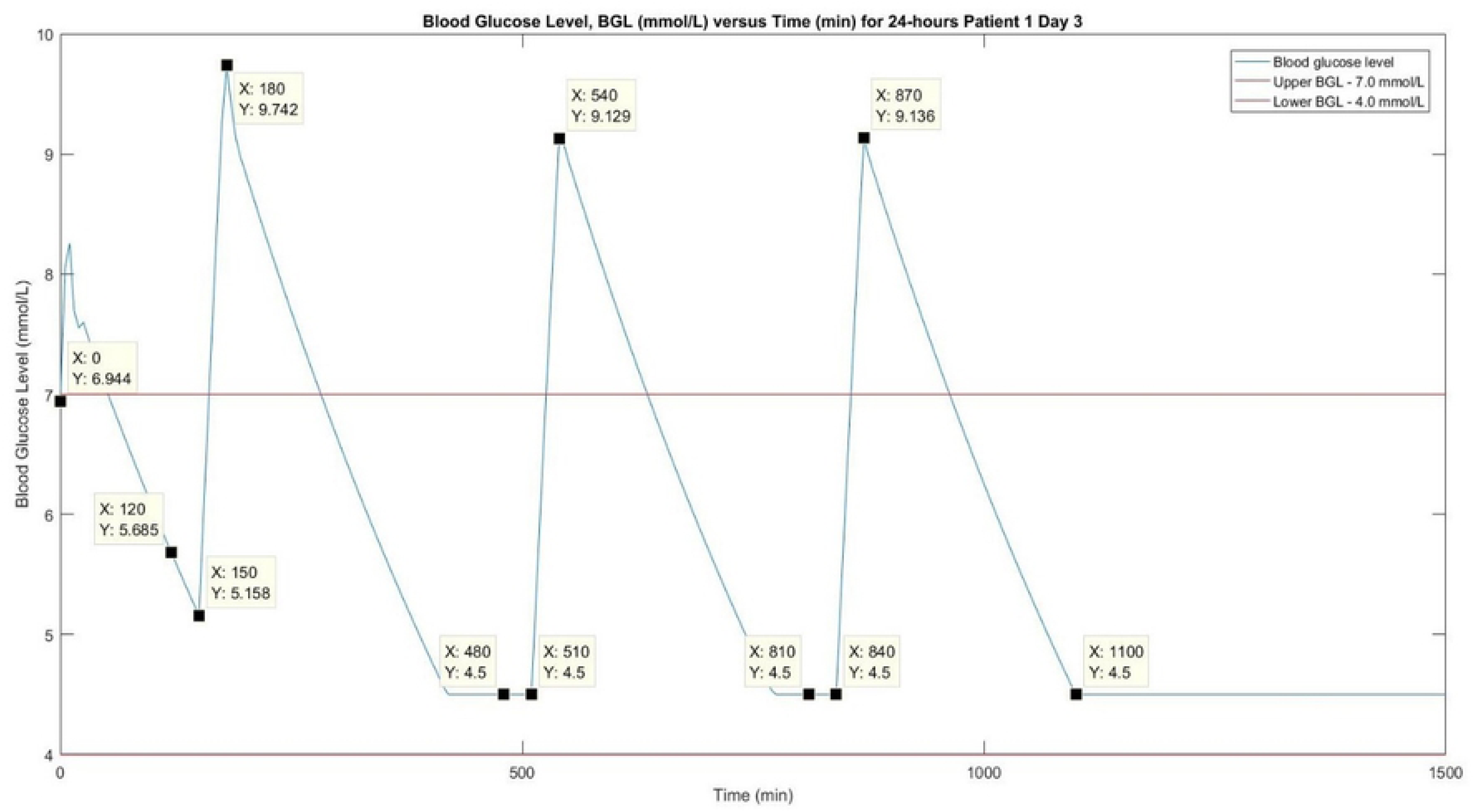
Simulation Work of Blood Glucose Level (mmol/L) versus.

**FIG. 14.**
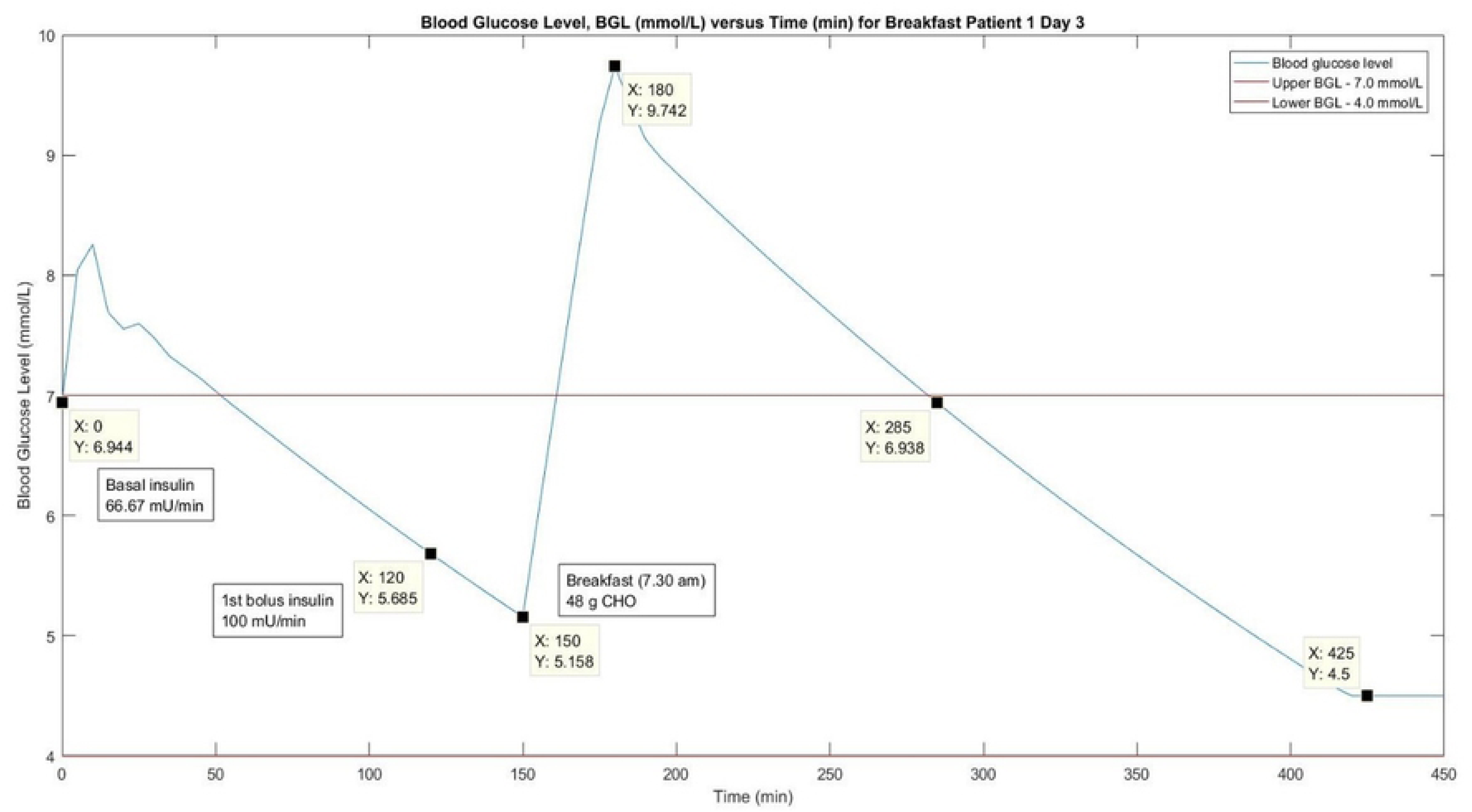
Simulation Work of Blood Glucose Level (mmol/L) versus.

**FIG. 15.**
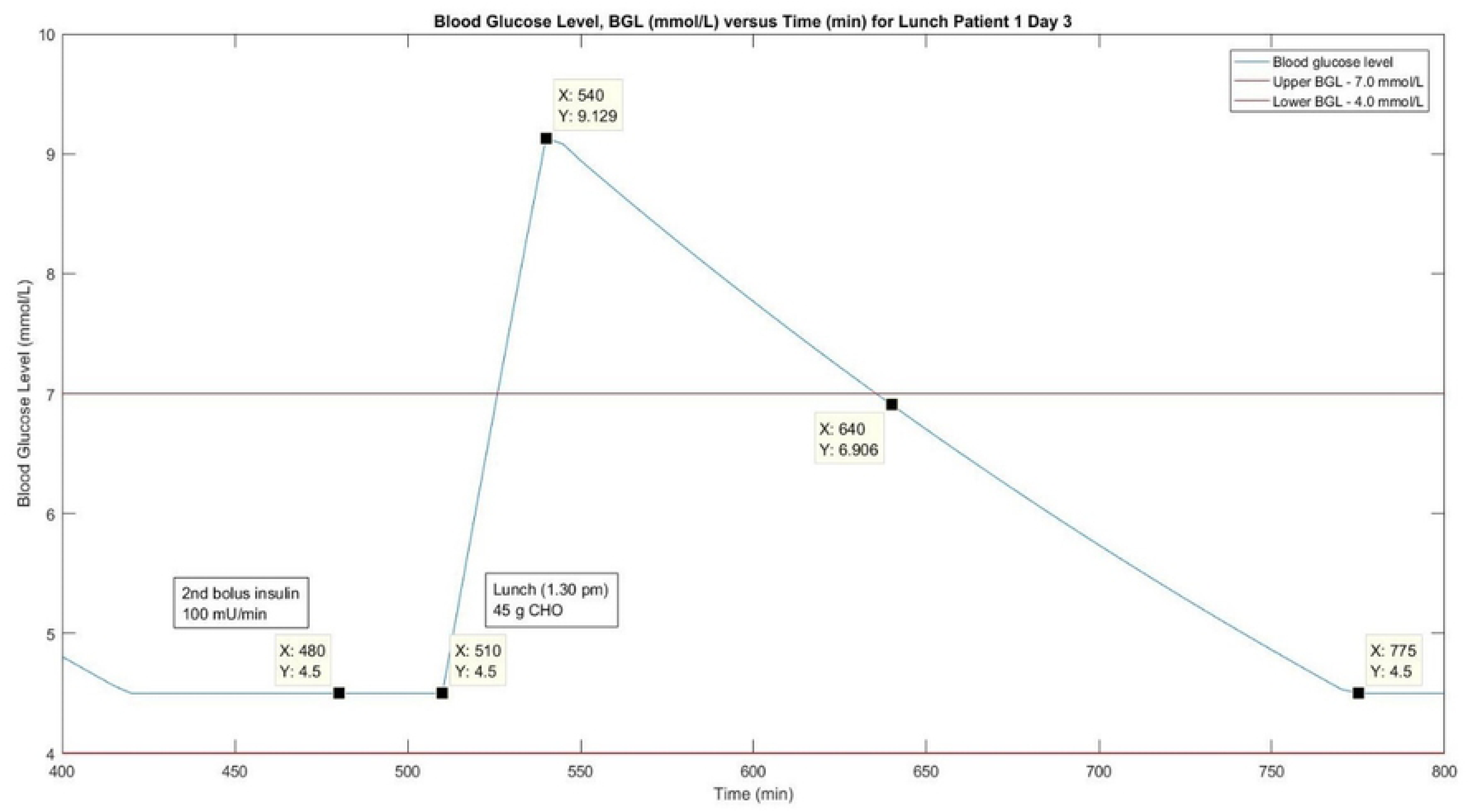
Simulation Work of Blood Glucose Level (mmol/L) versus.

**FIG. 16.**
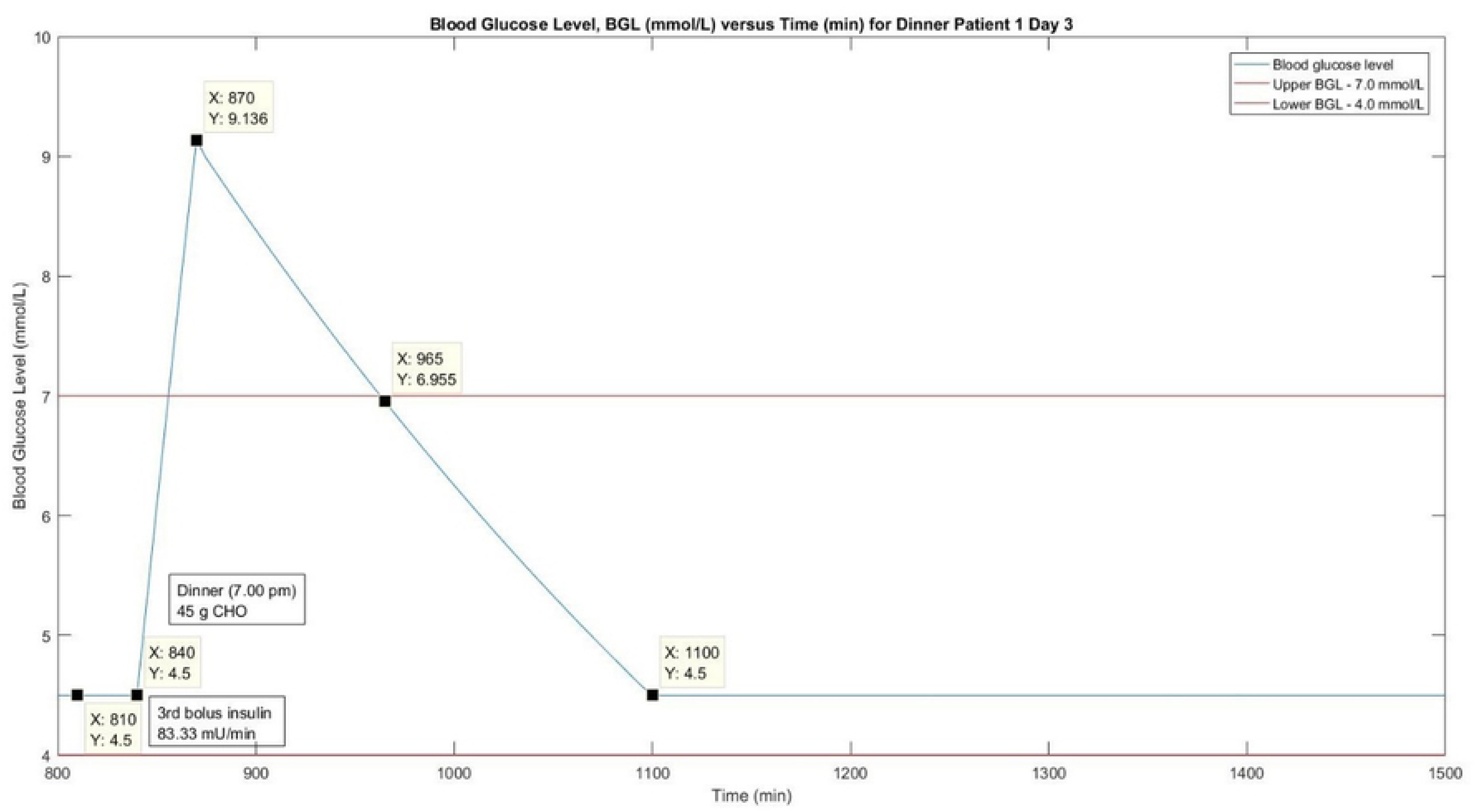
Simulation Work of Blood Glucose Level (mmol/L) versus.

Table 12 shows the comparison of peak BGL between clinical and simulation works on day 3. The BGL for clinical work was apparently higher than the simulation work. The highest BGL was recorded at 17.40 mmol/L during breakfast in the clinical work, while it was registered only at 9.74 mmol/L in the simulation. Table 13 shows that p value of x-variable 1 for clinical data was higher than 0.05 which indicated the data was not statistically reliable. This might be one of the reasons why the patient’s BGL was outside of the safe range most of the time in the clinical work. On the other hand, p values for simulation data were below 0.05 which indicated that the data were statistically reliable and very strong as shown in Table 14. It has proven again that simulation work is better at controlling BGL than the clinical work.

## Conclusions

In summary, this study had demonstrated that patient 1 was not able to maintain his BGL at safe range (4 to 7 mmol/L) most of the time within 24 hours for all three consecutive days as evidenced in the clinical works. Patient 1 often experienced hyperglycaemia; therefore, it was crucial to monitor his BGL so as to avoid long term complications in the future. The BGL recorded in the clinical works was not as reliable as in the simulation works. In clinical works, the patient needed to do SMBG before and two hours after every meal as well as carrying out MDI of insulins in order to maintain his BGL in the safe range. On the other hand, the BGL was continuously recorded and controlled on real time basis throughout the 24 hours duration in the simulation work. Furthermore, patient 1 was able to determine the appropriate bolus insulin needed and properly administered when there were fluctuations in BGL for future prediction purposes.

## Acknowledgements

The authors wish to acknowledge and extend their gratitude to the Malaysian Ministry of Higher Education (MOHE) and Research Management Centre (RMC), Universiti Teknologi MARA (UiTM) for the financial support (600-IRMI/DNA 5/3/BESTARI (060/2017)). The authors would also like to thank the Faculty of Chemical Engineering and UiTM Medical Specialist Centre, Faculty of Medicine, UiTM for all the support and assistance given in conducting this study.

## Author Disclosure Statement

Ayub Md Som and Nur’Amanina Mohd Sohadi wrote the paper; Nur’Amanina Mohd Sohadi, Noor Dyanna Andres Pacana and Nur Farhana Mohd Yusof participated in the programming works; Ayub Md Som, Noor Shafina Mohd Nor and Sherif Abdulbari Ali analysed the data collection and results; Noor Shafina Mohd Nor, Nur’Amanina Mohd Sohadi and Noor Dyanna Andres Pacana collected the blood sample results from the patient and conducted an interview session with the patient’s parent. All authors have no conflict of interest on this study.

